# PRMix: Primary Region Mix Augmentation and Benchmark Dataset for Precise Whole Mouse Brain Anatomical Delineation

**DOI:** 10.1101/2025.08.13.670045

**Authors:** Kunhao Yuan, Hanan Woods, Ülkü Günar, Digin Dominic, Ying Wu, Zhen Qiu, Seth G.N. Grant

## Abstract

The architecture of the mouse brain shares remarkable similarities with the human brain, making it an essential model for studying brain pathologies, synaptic diversity, and regional specialization. A key step in such studies involves registering molecular images to reference brain atlases, a process hindered by the difficulty of accurately delineating brain regions. Toward this, we have curated a collection of high-resolution, dual-fluorescence microscopy images, termed as dual-fluorescence mouse brain microscopy (DMBM) dataset, complemented by expert annotations of 118 subregions. This dataset provides unprecedented insights into the molecular and structural complexity of the mouse brain. However, its full potential for detailed whole-brain analysis is compromised by challenges such as boundary ambiguity and sample scarcity in existing automated segmentation methods, prompting the development of the primary region mix (PRMix) augmentation method. PRMix is specifically designed to expand these datasets, enhance the realism of synthetic data and minimize overlap between adjacent regions. Our approach, together with the curated dataset, achieves superior segmentation performance across the mouse brain compared with existing methods, setting a new benchmark in brain imaging research. Code and data are available at https://git-pages.ecdf.ed.ac.uk/dmbm-datasets-5c13cd/.

## 1. Introduction

The mouse brain is a fundamental model in neuroscience research owing to its structural and functional parallels with the human brain [33, 20]. Elucidating its molecular, synaptic, and cellular architecture is essential for understanding brain organization and function. Recent advances in molecular labeling, genetic tagging, and high-resolution imaging have enabled the generation of diverse, brain-wide datasets [37, 2]. However, the standard reference atlases that define brain region boundaries remain based on classical histology [14, 21], creating a critical need for accurate delineation methods that are compatible with modern molecular imaging modalities [37].

Manual delineation and annotation of brain regions are labor-intensive and prone to inter-annotator variability, limiting both reproducibility and scalability. Automated methods offer a scalable alternative, enabling highthroughput and standardized analyses. However, their performance critically depends on the availability of high-quality and diverse training datasets, which are particularly limited for modalities that integrate molecular information with anatomically accurate annotations. In this context, data augmentation plays a crucial role in enriching datasets by introducing variability and enhancing the robustness and generalization of automated models. Yet, most existing methods focus on a binary segmentation problem, such as tumor segmentation [17] or lesion segmentation [1], leaving more complex tasks, such as dense brain delineation, relatively underexplored. To address these challenges, we present the dual-fluorescence mouse brain microscopy (DMBM) dataset, a manually annotated collection of mouse brain sections that integrates both structural and molecular information. To improve the precision of automated brain region delineation, we propose primary region mix (PRMix), a novel augmentation method that preserves the original anatomical structure of the brain while minimizing regional overlaps, enabling realistic data synthesis and improving the precision of automated brain delineation [28].

## 2. Related work

### 2.1. Brain atlases

The Allen brain atlas [14] and Paxinos brain atlases [21] are widely used reference atlases that show delineated regions in planes of tissue sections. A core methodology used to create these atlases is Nissl staining [11], which highlights neuronal cell bodies and cytoarchitecture but lacks the resolution for molecular structures or synaptic connectivity. To overcome this limitation, fluorescent protein probes, such as those fused to endogenous synaptic proteins [37], self-labeling tags such as HaloTags [16], and antibody labeling [5], offer enhanced molecular insight and have been instrumental in developing single-synapse resolution maps of the mouse and human brain. In contrast to the cellular-level view of Nissl staining, protein-marked imaging provides superior subcellular detail.

More recently, advances in high-throughput 3D imaging have accelerated the creation of brain-wide datasets with cellular or subcellular resolution. Light-sheet fluorescence microscopy (LSFM), for instance, has been instrumental in generating whole-brain maps by enabling rapid optical sectioning of cleared tissue [22]. Similarly, serial two-photon tomography (STPT) has been employed to systematically image and reconstruct the entire mouse brain at micron-scale resolution, providing detailed cytoarchitectural and connectivity data [27].

While these methods provide invaluable insights into whole-brain architecture and function, they often rely on single molecular markers or are optimized for tracing long-range projections.

### 2.2. Dual-synaptic markers

Our work leverages a dual-synaptic marker strategy to dissect the molecular diversity of synapses. The two proteins we visualize, postsynaptic density protein 95 (PSD95) and synapse-associated protein 102 (SAP102), are key scaffolding molecules within the postsynaptic terminal of excitatory synapses but exhibit distinct expression patterns and functional roles throughout development [18] and across different brain regions [37, 4, 19, 6]. Imaging these molecules at single-synapse resolution across the brain has allowed for the generation of the first brain-wide synaptic maps in mammals across the lifespan [4, 2], in genetic models of neurodevelopmental disorders [37, 25] and in sleep deprivation [12].

The use of such multi-marker synaptic atlases has vast potential. They provide a crucial baseline for studying how synaptic composition is altered in models of disease, the effect of experience and learning, the impact of pharmacological interventions, and many other applications. Furthermore, this approach can be extended to include additional synaptic markers to create even more detailed synaptome maps [37], enabling researchers to undertake advanced functional [26] and connectomic [31] studies.

### 2.3. Image augmentation

Deep learning-based image analysis, especially in medical imaging, often faces data scarcity [15, 7]. To mitigate this, image augmentation techniques, which were originally popularized in natural image recognition, have been widely adopted. Traditional approaches apply geometric transformations such as flipping and affine adjustments (rotation, scaling, translation) to teach models spatial invariance, as well as intensity perturbations (brightness and contrast shifts) to ensure robustness to variations in imaging conditions [13, 24, 3].

Building on these foundational methods, more recent work has explored mixing-based augmentations. MixUp [34] introduced linear image combinations to expand datasets and regularize training, and was later extended to segmentation tasks [8]. Recent advances include patch-based augmentation [32], scribble-based methods for medical imaging [35], semantic-aware augmentation [36, 29], and self-adaptive blending to address background inconsistencies [38]. Although existing mixing methods can improve performance on object-centric tasks, they often fail to preserve the global anatomical topology required for whole-brain delineation. PRMix, with its primary region sampling and overlap-aware augmentation, is specifically designed to address this gap.

## 3. The dual-fluorescence mouse brain microscopy dataset

### 3.1. Data collection

The dataset includes brain sections from 96 mice (48 male, 48 female), aged 1–12 months. Fifty-one were C57BL/6 × 129S5 crosses expressing *Psd95*^*eGFP*^, *Sap102*^*mKO2*^, and *GluN1*, while 45 were C57BL/6 expressing *Psd95*^*eGFP*^ and *Sap102*^*mKO2*^. Mice were perfused with sodium pentobarbital and saline-PFA. Brains were dissected, post-fixed, and cryo-embedded in OCT compound. Parasagittal sections (18 *μ*m) were cut referencing Allen

### 3.2. Dataset statistics

The collected dataset has **102** dual-fluorescence whole-brain images (*n* = 96), with a median resolution of 6383×12531, encompassing **118** well-defined anatomical subregions from the sagittal mouse brain slices. Three exemplar images are shown in Figure 2. The total number of pixels for each subregion is log-rescaled and summarized in Figure 1a, which highlights significant variations in pixel counts across subregions. Notably, the foregroundto-background ratio is approximately **0.84**, making the proposed dataset highly challenging while informative for neuroscience and medical research. mouse brain atlas slices 11–12 [14]. Whole-brain imaging was performed on a Nikon Eclipse Ti2 with a spinning disk confocal system, and 856×812 individual tiles (per image) were stitched into full images with 16× downsampling. Subregions were manually delineated in ImageJ using PSD95/SAP102 protein markers, guided by the Allen brain atlas.

**Figure 1.**
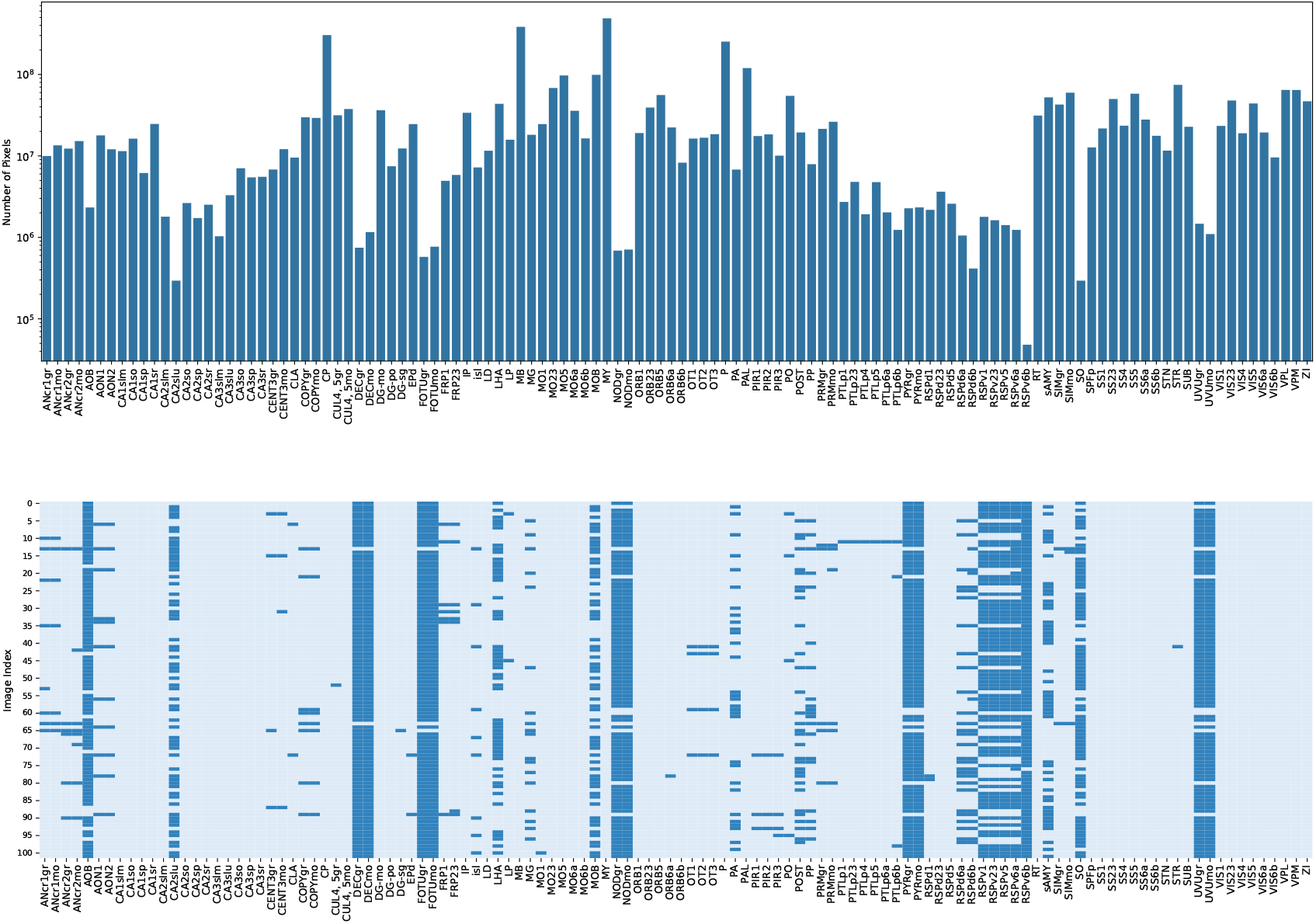
(Top) a. Distribution of pixel counts across regions. (Bottom) b. Region availability per image with light blue indicating presence and dark blue representing absence.

**Figure 2.**
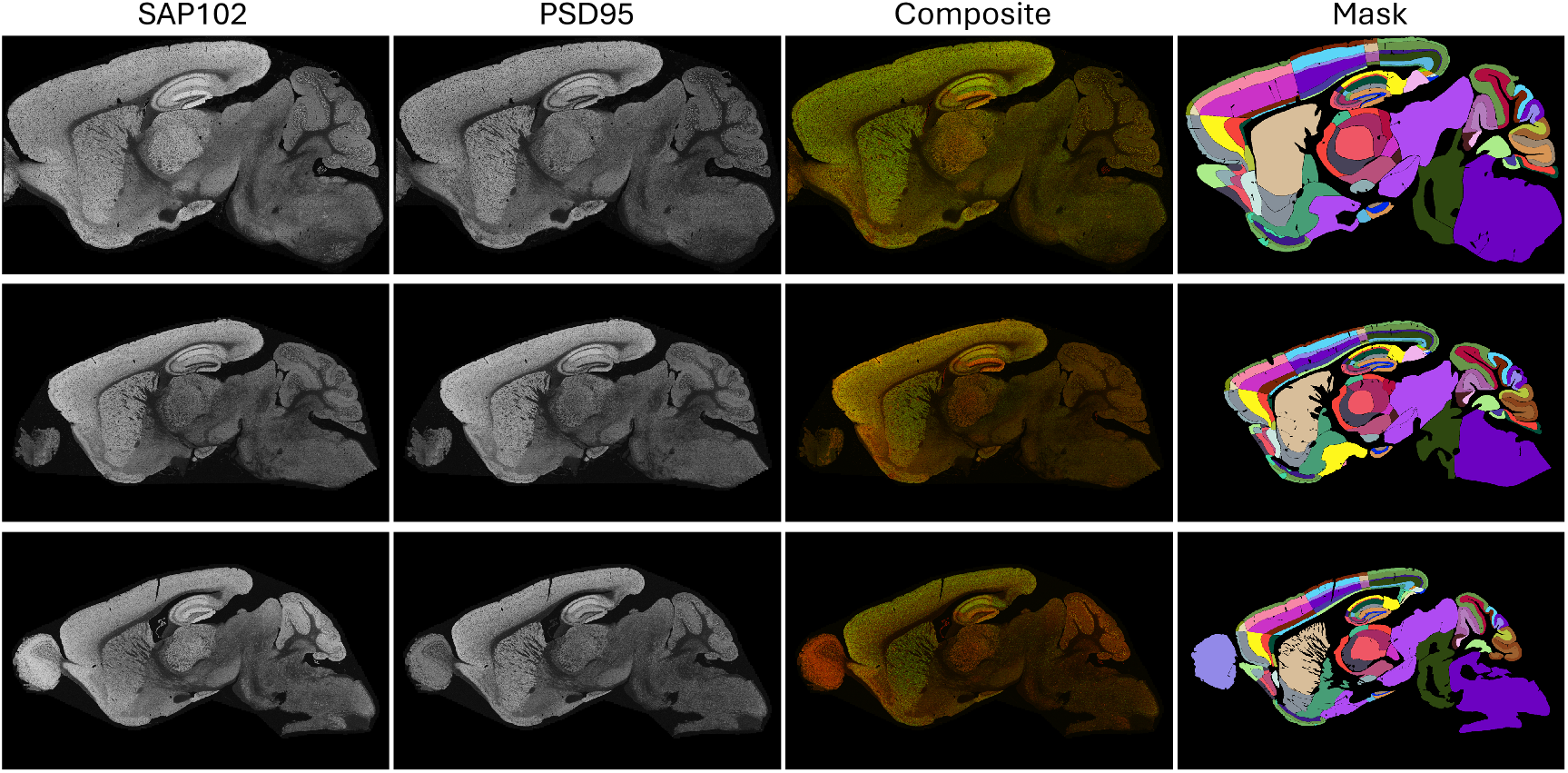
Exemplar images from the curated dataset. From left to right: whole-brain images stained for SAP102 and PSD95, the dual-channel composite image (SAP102 in red, PSD95 in green), and the corresponding manually annotated brain regions.

Figure 1b illustrates the missing subregions for specific image IDs. Due to the location of the slices during sectioning, certain areas, such as AOB and RSPv6b, are present in only two slices and are considered less reliable and representative. We thus excluded them in evaluation but kept the information in the figure for completeness.

## 4. Methods

### 4.1. Motivation

While we introduced a high-resolution, dual-fluorescence dataset, microscopy data are scarcer than for CT [30] or MRI [17] due to their extremely large size (e.g., 102k×200k pixels) and low throughput, and thus insufficiently diverse on its own to eliminate augmentation. This scarcity, coupled with biological variability, makes augmentation essential. PRMix is thus motivated to enhance data diversity and realism for effective modelling. The overall diagram of the proposed PRMix is illustrated in Figure 3, with the subsequent paragraphs detailing its design objectives and implementation.

**Figure 3.**
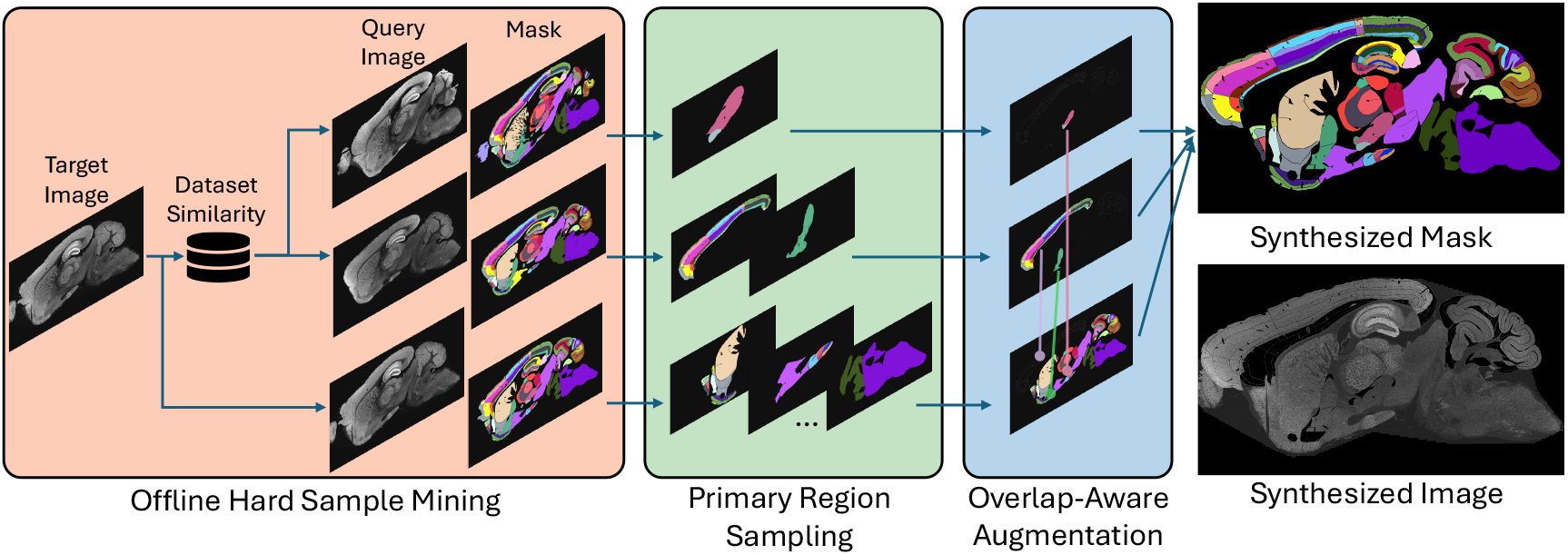
Overview of the proposed PRMix, consisting of three modules (1) offline hard-sample mining (HSM), (2) primary region sampling (PRS) and (3) overlap-aware augmentation (OAA). Only a single fluorescent marker is shown for clarity.

### 4.2. Offline hard-sample mining

According to our initial experiments, random sample mixing can introduce feature inconsistencies, while exclusively selecting visually similar samples lacks sufficient challenge for effective learning. We therefore designed a strategy to ensure each target image is paired with multiple query images from a diverse range of pre-computed similarities across the whole dataset, termed as offline hard-sample mining. To minimize pixel-level noise while preserving semantic meaning, we compute similarity scores by summing over all pixel locations where the query mask matches the target mask, followed by normalization by the number of foreground pixels in the target mask. For each target image, similarity scores are computed against the remaining training images and sorted. The top 20% most similar samples are designated as “easy samples”, while the remaining 80% are labeled as “hard samples”. During data augmentation, we mix “easy” and “hard” samples in varying proportions, aiming to find a balance between challenge and learnability. Empirically, oversampling 80% from the hard-sample set yielded the best results. Details of the different sampling ratios are summarized in Table 5a.

### 4.3. Primary region sampling

Based on anatomical and functional criteria [14, 20], we grouped the 118 brain subregions into 11 primary regions: cerebellum (CB), thalamus (TH), midbrain (MB), hindbrain (HB), isocortex (ISO), hypothalamus (HY), olfactory areas (OLF), cortical subplate (CTXsp), striatum (STR), pallidum (PAL), and hippocampal formation (HPF), as illustrated in Figure 4. These primary regions serve as the fundamental units for mixing. During mixing, we only sampled 3–9 of these regions from multiple sources to replace those in target image, boosting diversity while preserving feature consistency. The primary region sampling offers two key advantages: (1) preserving the anatomical structure of the brain image and (2) avoiding potential subregional overlaps during the mixing process. More importantly, our design incorporates mixing at an intermediate semantic level, setting it apart from existing object-centric approaches [34, 32, 36, 29]. This makes it particularly well-suited for dense foreground tasks such as whole-brain delineation.

**Figure 4.**
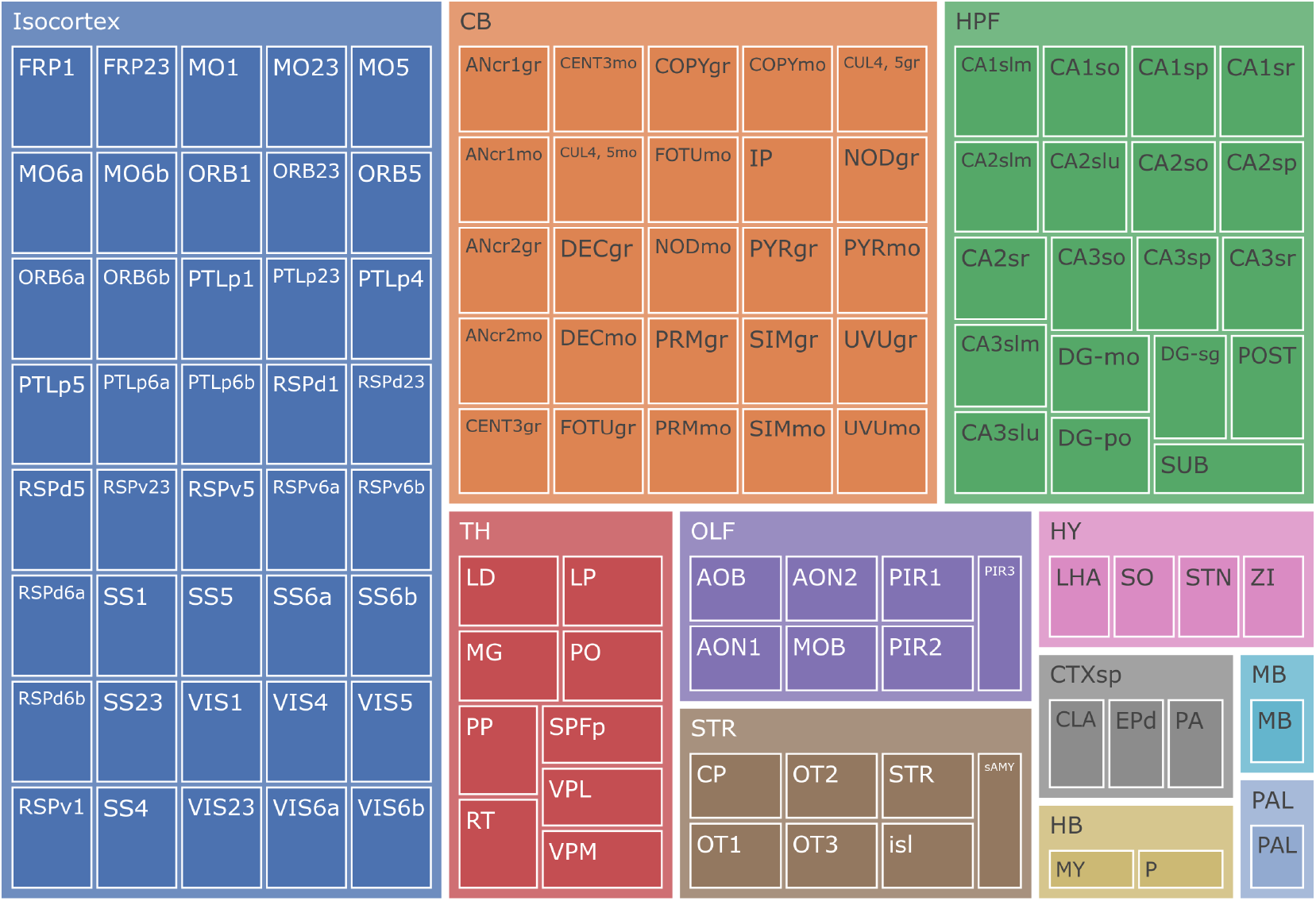
Illustration of the 11 defined primary regions and the corresponding affiliations of the 118 subregions.

### 4.4. Overlap-aware augmentation

To ensure those sampled regions align in location and size with the target image, we implemented an overlap-aware augmentation module. This module begins by applying random affine transformations— such as rescaling, rotation, and translation— to the query masks for individual primary regions. Corresponding regions are then cut out from the target masks, leaving behind a complementary mask which is used to calculate intersections with the query mask. If the intersection is below a certain threshold *τ*, the query region will be pasted directly onto the complementary masks at its original location. Otherwise, a greedy search for optimal affine parameters will begin to ensure a minimum intersection where possible. Once the greedy search converges, the optimized affine transformations are reapplied to the query images and masks, followed by the final pasting process. The overall process is outlined as pseudo-code in Algorithm 1.

#### Algorithm 1

OAA: Overlap-Aware Augmentation

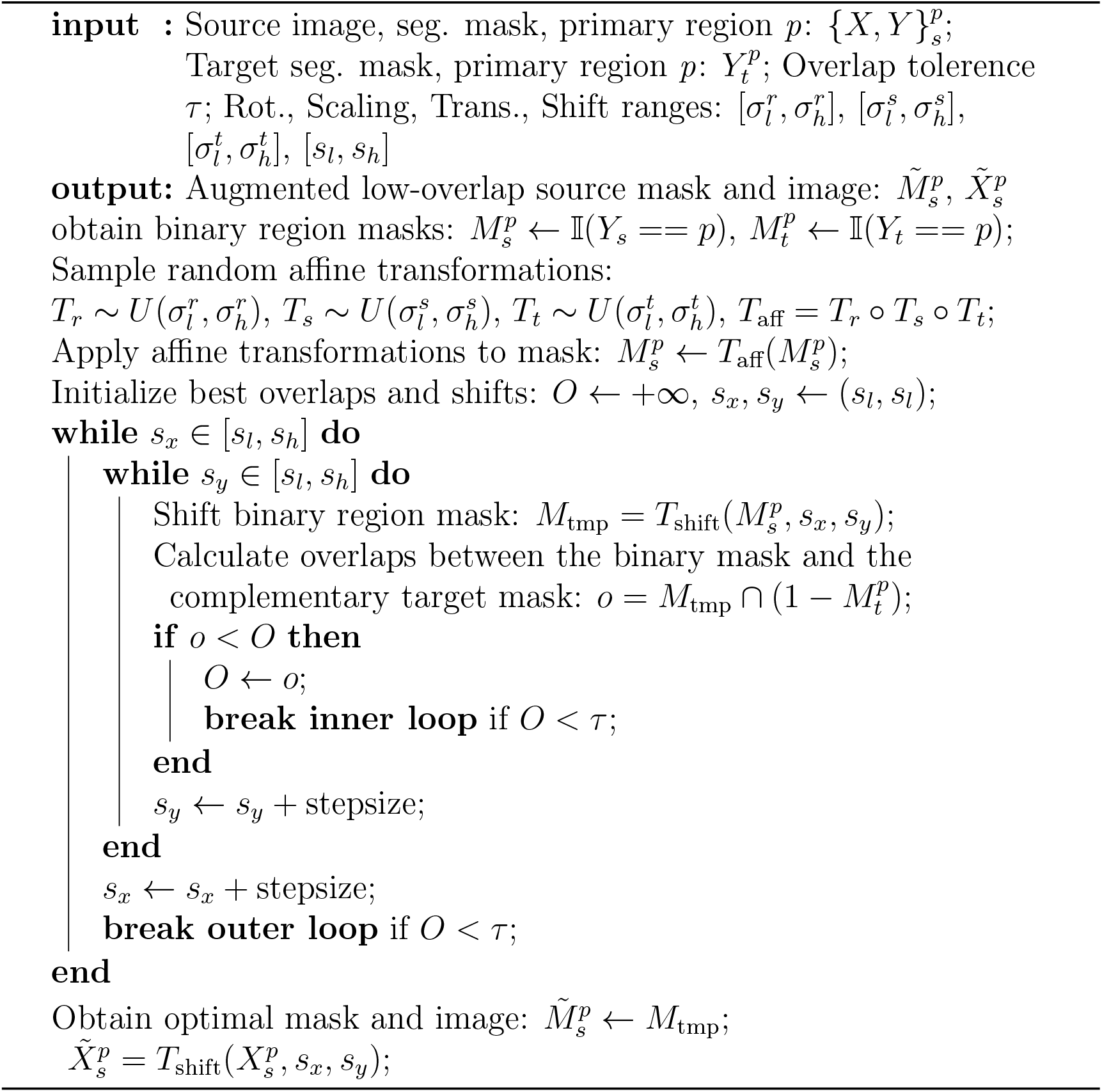

### 4.5. Comparison with existing mixing approaches

Before formalizing the proposed PRMix, we compare it with existing mixing-based augmentations. Starting from the simplest MixUp [34], it blends a pair of images, *i.e*. source image *X*_*s*_ and target image *X*_*t*_ and their corresponding labels *Y*_*s*_ and *Y*_*t*_. The synthesized image and label are generated through:

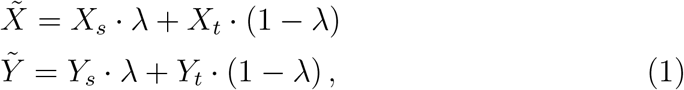

where *λ* ~ *U* (0, 1) is a controlling factor. For CutMix [32] and CarveMix [36], their mixing strategies can be summarized as:

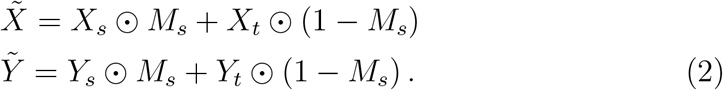

⊙ is the element-wise dot product and *M*_*s*_ represents a binary mask used to sample areas from the source image, which can be either rectangular regions (CutMix) or semantic regions (CarveMix). Taking both source image and target image semantics into account, we yielded the basic form of our PRMix:

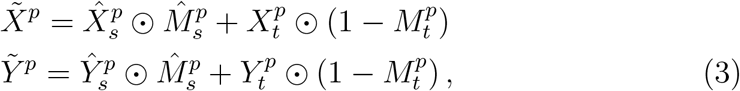

where 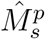 is the overlap-mitigated semantic mask for primary region *p*, retrieved with 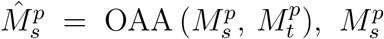 is the union of total *k* subregions within a primary region, known as 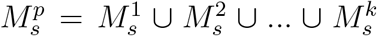, and the adapted source image 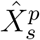 is generated with the overlap-mitigated semantic mask, using 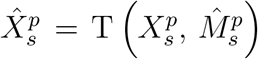. Additionally, to enhance the versatility and variability of the dataset, we further extended the proposed PRMix to multiple source images, simply by denoting the above process as 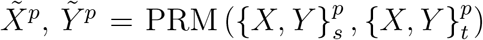, and iterate the process multiple times via:

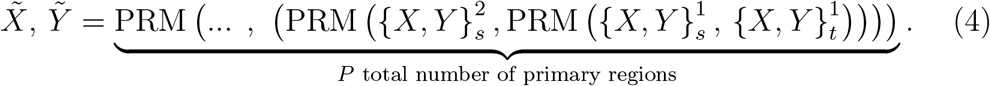

This enables mixing at an intermediate semantic level and allows generalization beyond a single image pair, unlike traditional augmentation methods where repeated lower-level mixing often leads to occlusion and ambiguity.

Exemplar synthesized images from different methods are illustrated in Figure 5. MixUp [34] generates ambiguous images, whereas CutMix [32] and CarveMix [36] focus on single regions and struggle to preserve global anatomy. By contrast, our method produces the most realistic and diverselooking images.

**Figure 5.**
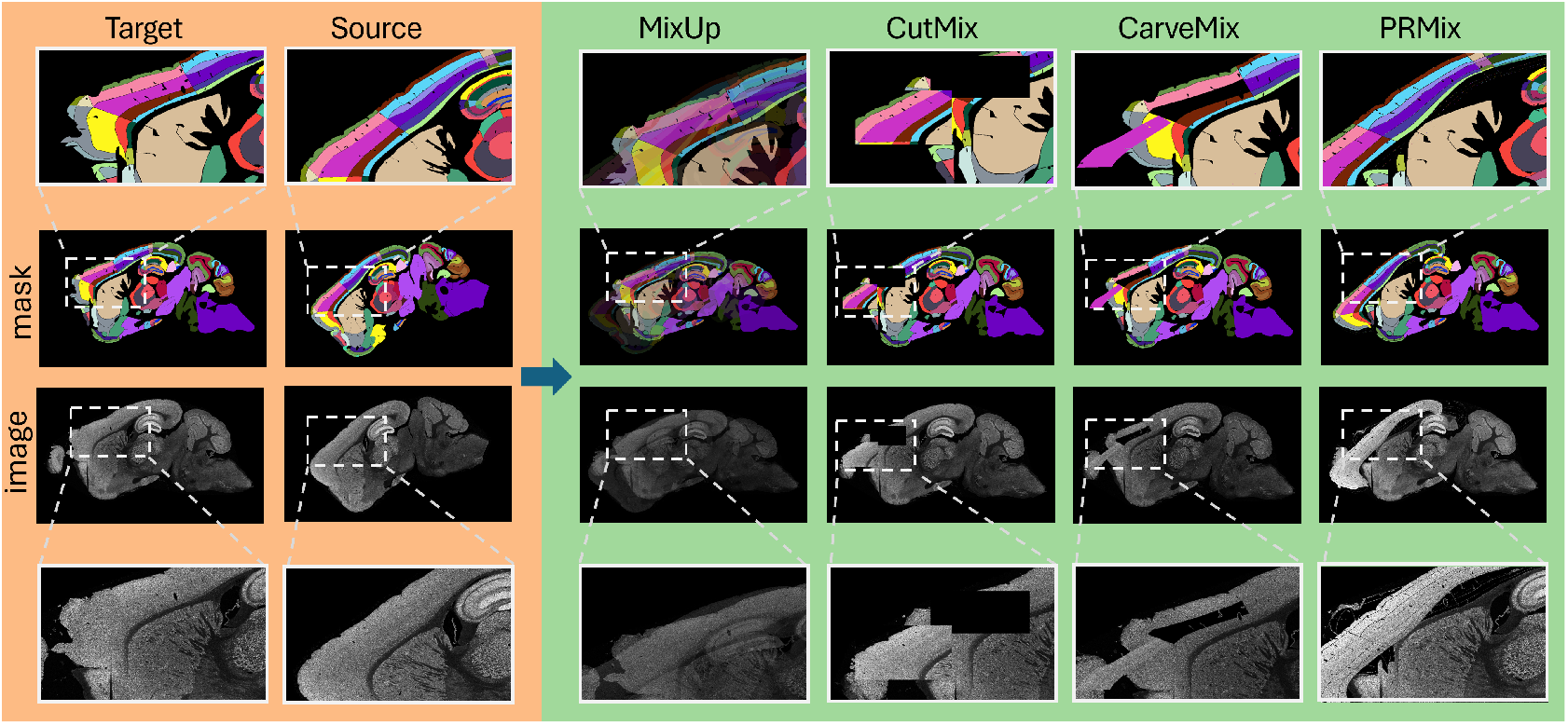
The exemplar results from different methods. The middle two rows show the original image and its corresponding label mask, while the top and bottom rows provide zoomed-in views of the manipulated regions. Best viewed in color.

## 5. Experiments

### 5.1. Implementation details

To ensure a fair comparison between different augmentation methods and minimize the influence of model architectures, all experiments were conducted using the state-of-the-art medical image segmentation model nnUNet [9, 10], which enhances the classic UNet [23] with task-specific configurations. The proposed PRMix was executed offline to generate an augmented dataset with different folds, optimizing training efficiency. In addition, standard preprocessing techniques, including default intensity normalization, foreground oversampling, color jittering, and affine transformations, were applied for all methods during training. The model was trained with an input patch size of 768 × 1536, approximately 1*/*64 of the median image size, using 2 patches per GPU and 250 minibatches per epoch. The collected dataset was split into 80 training/validation and 22 strictly isolated and unaugmented testing samples, and all experiments were conducted using 5-fold cross-validation on two NVIDIA RTX 4090 GPUs.

The augmented training sets were generated with varying folds, namely 5 × (~ 400 images), 10× (~ 800 images) and 20 × (~ 1600 images), by excluding cases where source regions are absent from the target image. Due to the large size of the whole-brain image, patch-wise training has been employed. Under the given patch size and batch settings, the model requires approximately 2,500 iterations to process the entire training dataset once—a process that takes about two days on a single GPU. Consequently, the baseline model was trained for 12 full epochs as a reference, while the proposed PRMix model was evaluated with 3, 6, and 12 full epochs for comparison. Hereafter, unless otherwise specified, all mentions of “epochs” refer to full epochs.

### 5.2. Experimental results

We compared PRMix with existing methods in Table 1, evaluating categorical average precision, recall, F1 and mIoU. All results are averaged over 5-fold runs unless stated otherwise. The best score is highlighted in dark gray and the second best is in light gray. Although MixUp [34] utilizes an augmented dataset, it introduces ambiguity in dense segmentation, leading to the lowest precision and recall among all augmentation strategies. While CutMix [32] and CarveMix [36] improve F1 and mIoU over the baseline, they fail to capture all true pixels, resulting in similar performance to MixUp. We hypothesize that this occurs because these methods generate synthetic samples without considering spatial locations, leading to overlapping regions that make distinguishing individual objects more challenging. By contrast, our proposed PRMix significantly outperforms all compared methods across all metrics, achieving the highest F1 (74.66) and mIoU (61.78) among all methods.

**Table 1:** Results on the isolated test set using different mixing strategies, which are obtained with a 5-fold average. ±: standard deviations over 5 runs.

| Method | Precision | Recall | F1 | mIoU |
| --- | --- | --- | --- | --- |
| Baseline | 66.35±0.87 | 66.55±0.40 | 65.33±0.49 | 54.33±0.48 |
| MixUp 20× [34] | 74.71±0.43 | 73.77±0.39 | 73.30±0.53 | 60.73±0.40 |
| CutMix 20× [32] | 75.05±0.42 | 73.83±1.02 | 72.97±0.79 | 60.67±0.62 |
| CarveMix 20× [36] | 74.10±1.07 | 74.44±0.49 | 73.30±0.85 | 60.87±0.56 |
| PRMix 20× | 75.23±0.57 | 75.31±0.33 | 74.66±0.29 | 61.78±0.33 |

Qualitative results from different methods are illustrated in Figure 6. The top two rows show under-represented regions, such as the UVUgr (primary region: CB), which are missing in the results of Baseline, CutMix, and CarveMix. The mispredictions by CutMix and CarveMix are likely due to severe overlaps introduced during mixing, while the Baseline model’s failure can be attributed to insufficient training data for this small region. The middle three rows demonstrate the challenge of distinguishing background and dark regions, where most methods struggle. By contrast, the proposed PRMix produces results that align more closely with manual annotations. The last row presents an extremely ambiguous sample, where the MOB region (primary region: OLF) appears damaged—likely due to sectioning artifacts. The corresponding manual annotation reflects the expert’s decision to exclude the damaged area. While none of the tested methods fully eliminate this ambiguous region, PRMix exhibits the minimum discrepancy.

**Figure 6.**
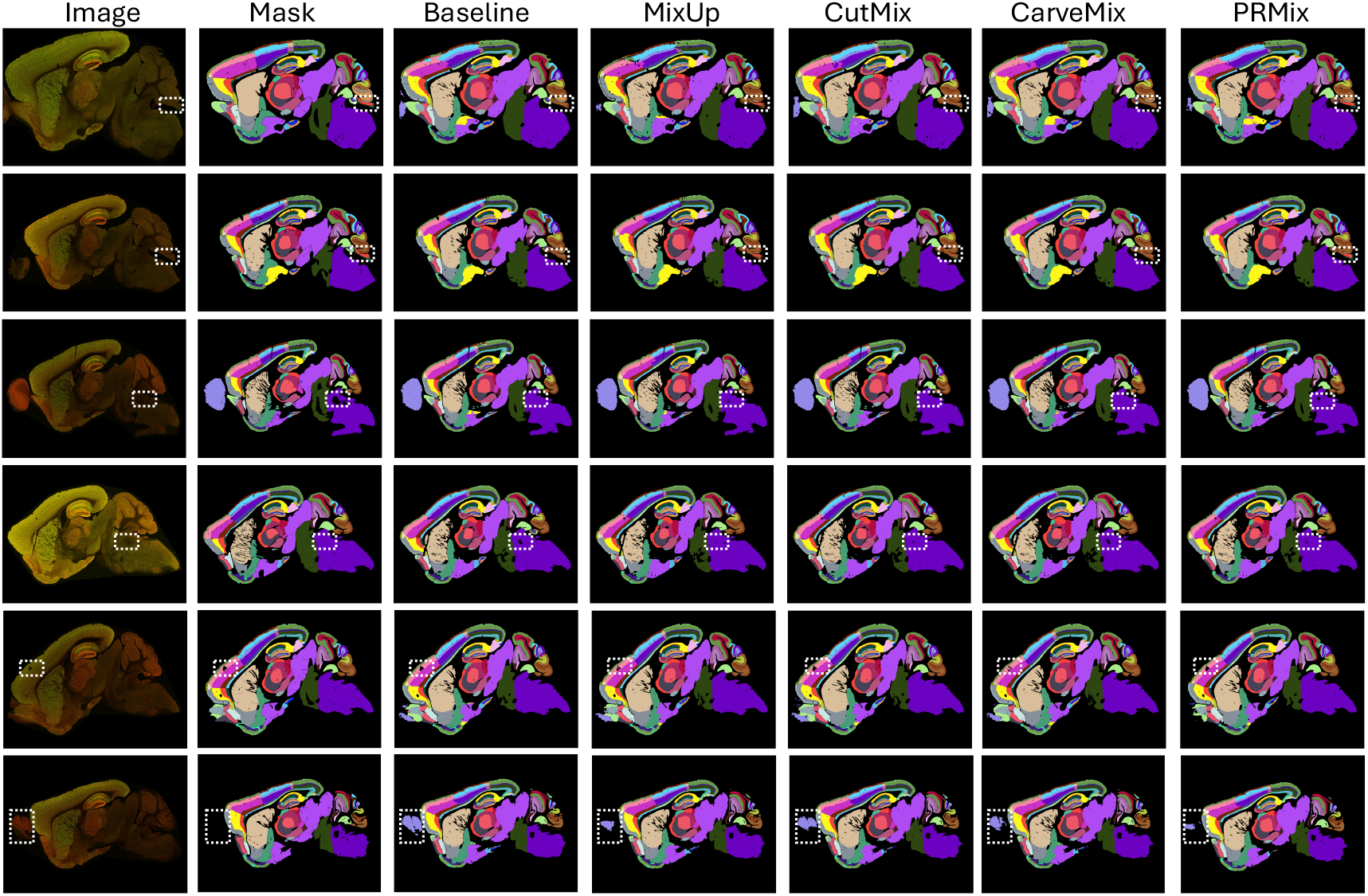
Visualization of results from different methods on randomly selected testing samples. Dashed white rectangles indicate regions with the most significant discrepancies between methods. Best viewed digitally.

While qualitative comparisons can highlight differences between methods, they are susceptible to selective bias. To mitigate this, we conducted a rigorous statistical analysis using a paired t-test to determine the significance of the observed differences.

For this analysis, we first calculated the F1 and mIoU scores for every sample in the test set across all methods and training folds. The scores were then grouped, pairing the reference method’s results with those of each comparator method. Each pair of groups was then subjected to a paired t-test, with the results detailed in Table 2. When using the baseline as the reference method, PRMix and all other methods significantly outperformed it. When compared against existing methods, PRMix still showed statistically significant improvements over MixUp and CarveMix. Although PRMix did not achieve significance against CutMix, the p-values for F1 and mIoU were 7.89e-2 and 5.65e-2, respectively—both close to the significance threshold.

**Table 2:**
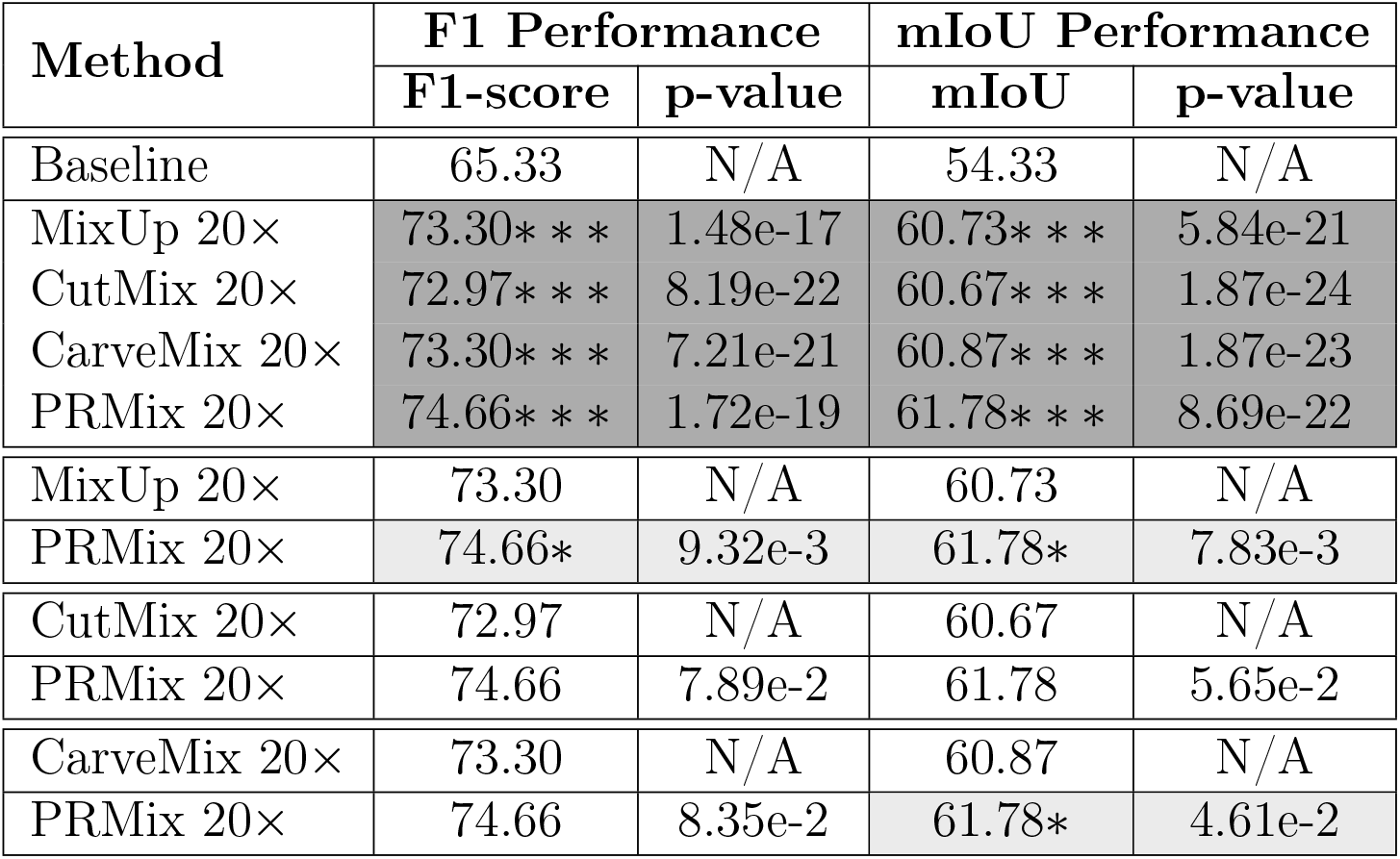
Paired t-test p-values for F1 and mIoU, computed on test samples. *: P < 0.05, **: P < 0.01 and ***: P < 0.001. One-sided tests evaluate whether the selected method yields significantly higher F1 or mIoU than the reference. In each subgrid, the method above the horizontal line is the reference and the one below is the tested method.

| Method | F1 Performance |  | mIoU Performance |  |
| --- | --- | --- | --- | --- |
|  | F1-score | p-value | mIoU | p-value |
| Baseline | 65.33 | N/A | 54.33 | N/A |
| MixUp 20× | 73.30*** | 1.48e-17 | 60.73*** | 5.84e-21 |
| CutMix 20× | 72.97*** | 8.19e-22 | 60.67*** | 1.87e-24 |
| CarveMix 20× | 73.30*** | 7.21e-21 | 60.87*** | 1.87e-23 |
| PRMix 20× | 74.66*** | 1.72e-19 | 61.78*** | 8.69e-22 |
| MixUp 20× | 73.30 | N/A | 60.73 | N/A |
| PRMix 20× | 74.66* | 9.32e-3 | 61.78* | 7.83e-3 |
| CutMix 20× | 72.97 | N/A | 60.67 | N/A |
| PRMix 20× | 74.66 | 7.89e-2 | 61.78 | 5.65e-2 |
| CarveMix 20× | 73.30 | N/A | 60.87 | N/A |
| PRMix 20× | 74.66 | 8.35e-2 | 61.78* | 4.61e-2 |

### 5.3. Ablations

We performed a comprehensive ablation study to investigate the effects of several key factors: dataset scale, training duration, the number of mixed images, the fluorescent channels used, the proportion of hard samples, and the contribution of each proposed module. The results are detailed in Table 3, 4 and 5.

**Table 3:** Ablation study on the impact of dataset scaling in augmented folds and training duration.

| (a) Scaling effect |  |  |  | (b) The number of training epochs |  |  |  |
| --- | --- | --- | --- | --- | --- | --- | --- |
| Method | Epochs | F1 | mIoU | Method | Epochs | F1 | mIoU |
| Baseline | 12 | 65.33 | 54.33 | Baseline | 12 | 65.33 | 54.33 |
| PRMix 5× | 12 | 69.82 | 58.14 | PRMix 20× | 3 | 68.53 | 56.75 |
| PRMix 10× | 12 | 70.05 | 58.13 | PRMix 20× | 6 | 69.92 | 57.97 |
| PRMix 20× | 12 | 74.66 | 61.78 | PRMix 20× | 12 | 74.66 | 61.78 |

**Table 4:** Ablation on the number of mixing images and fluorescent channels.

| (a) The number of mixing images |  |  | (b) Fluorescence channels |  |  |
| --- | --- | --- | --- | --- | --- |
| Method | F1 | mIoU | Method | F1 | mIoU |
| Baseline | 65.33 | 54.33 | Baseline | 65.33 | 54.33 |
| PRMix Dual | 74.34 | 61.58 | PRMix_SAP102 | 69.92 | 57.77 |
| PRMix Tri | 74.66 | 61.78 | PRMix_PSD95 | 70.00 | 58.44 |
| PRMix Quad | 71.03 | 59.50 | PRMix | 74.66 | 61.78 |

**Table 5:** Ablation on the portion of hard samples and key components.

| (a) The portion of hard samples in HSM |  |  | (b) Key components |  |  |  |  |
| --- | --- | --- | --- | --- | --- | --- | --- |
| Method | F1 | mIoU | PRS | HSM | OAA | F1 | mIoU |
| Baseline | 65.33 | 54.33 | ✓ |  |  | 73.70 | 60.76 |
| PRMix 20% | 72.73 | 59.98 | ✓ | ✓ |  | 71.17 | 59.21 |
| PRMix 50% | 69.71 | 58.02 | ✓ |  | ✓ | 70.60 | 58.57 |
| PRMix 80% | 74.66 | 61.78 | ✓ | ✓ | ✓ | 74.66 | 61.78 |

The results in Table 3(a) highlight the benefit of extensive data augmentation. Performance steadily improves as augmentation increases from 5× to 20×, where the model achieves its highest F1 and mIoU scores, surpassing the baseline by a large margin. Table 3(b) confirms that performance also correlates with longer training. However, comparing the two factors reveals that augmentation provides a greater advantage than training duration. For example, PRMix 20× with just 3 epochs of training outperforms PRMix 5× with 12 epochs, demonstrating that exposure to more diverse samples is more critical than longer training with less diverse data.

Our investigation into the number of mixed images (Table 4(a)) indicates that three is the optimal number. Mixing two images provides minimal variation and risks overfitting, whereas mixing four disrupts the target image’s structure. Empirically, mixing three images achieves the best balance and the highest performance. In Table 4(b), we assess the impact of fluorescence channels. The dual-channel model significantly outperforms models trained on either the SAP102 or PSD95 channel alone, validating our methodological design. We also note that the PSD95 model slightly outperforms the SAP102 model, likely because its clearer expression in maturer synapses [18] helps in distinguishing regional boundaries.

Table 5a shows that overemphasis on “easy” samples (the 20% “hard” setting) leads to suboptimal performance, likely due to insufficient feature diversity. In addition, the 50%:50% configuration does introduce greater diversity but still lacks enough hard examples to support learning a more generalizable representation, which accounts for its performance gap compared with the 80%–hard setting. Table 5b shows that integrating primary region sampling (PRS) alone yields a noticeable performance improvement compared to the baseline. However, further adding either HSM or OAA individually leads to performance degradation, suggesting that the “easy” setting alone does not benefit from OAA, and that introducing additional hard samples through HSM increases task complexity. In contrast, combining both HSM and OAA produces a substantial performance gain, underscoring their synergistic effect in enhancing the dataset quality.

## 6. Conclusion

We curated the DMBM dataset, a high-resolution, expert-annotated collection capturing 118 brain subregions with unprecedented molecular and structural detail of the synaptic organization. For the first time, we demonstrate that integrating dual-fluorescence synaptic markers enables this dataset to serve as a comprehensive benchmark for regional delineation of the mouse brain.

To further improve the accuracy of automated segmentation methods, we introduced PRMix, a novel data augmentation that enables realistic data synthesis while preserving anatomical structures, achieving fine-grained brain region delineation beyond existing methods. By combining the DMBM dataset with PRMix augmentation, our work sets a new standard for mouse brain delineation and provides a robust framework for advancing neuroscience and biomedical imaging.

The resulting automated delineator offers a >10-fold reduction in processing time, resolving boundary ambiguities and minimizing the inconsistencies of manual annotation. This efficiency dramatically lowers the barrier to creating large-scale, high-accuracy atlases. Looking forward, the adaptability of our framework opens critical new research avenues. It enables the creation of dynamic atlases that chart brain development or disease progression and facilitates high-throughput screening of anatomical changes in preclinical models. Furthermore, its potential to generalize across molecular markers and species provides a powerful tool for comparative neuroanatomy and for integrating imaging data with other modalities, such as spatial transcriptomics and connectomics, paving the way for a more holistic understanding of brain architecture.

## Data and code availability

The data and code used for this study have been made public at https://git-pages.ecdf.ed.ac.uk/dmbm-datasets-5c13cd/. For the purpose of open access, the author has applied a CC-BY public copyright license to any Author Accepted Manuscript version arising from this submission.

## CRediT authorship contribution statement

**Kunhao Yuan**: Data curation, Conceptualization, Methodology, Writing - original draft, Writing - reviewing & editing. **Hanan Woods**: Data collection, Data curation, Writing - review & editing. **Ülkü Günar**: Data collection, Data curation. **Digin Dominic**: Visualization, Resources. **Ying Wu**: Formal analysis. **Zhen Qiu**: Formal analysis, Writing - review & editing. **Seth G.N. Grant**: Conceptualization, Writing – review & editing, Resources, Supervision, Funding acquisition.

## Acknowledgment

This work was funded by the Wellcome Trust (302077/Z/23/Z, 218293/Z/19/Z, 221295/Z/20/Z) and Simons Initiative for the Developing Brain (SIDB) under the Simons Foundation for Autism Research Initiative (529085). We thank C. Davey for editing.

